# RetroPath2.0: a retrosynthesis workflow for metabolic engineers

**DOI:** 10.1101/141721

**Authors:** Baudoin Delépine, Thomas Duigou, Pablo Carbonell, Jean-Loup Faulon

## Abstract

Synthetic biology applied to industrial biotechnology is transforming the way we produce chemicals. However, despite advances in the scale and scope of metabolic engineering, the bioproduction process still remains costly. In order to expand the chemical repertoire for the production of next generation compounds, a major engineering biology effort is required in the development of novel design tools that target chemical diversity through rapid and predictable protocols. Addressing that goal involves retrosynthesis approaches that explore the chemical biosynthetic space. However, the complexity associated with the large combinatorial retrosynthesis design space has often been recognized as the main challenge hindering the approach. Here, we provide RetroPath2.0, an automated open source workflow for retrosynthesis based on generalized reaction rules that perform the retrosynthesis search from chassis to target through an efficient and well-controlled protocol. Its easiness of use and the versatility of its applications make of this tool a valuable addition into the biological engineer bench desk. We show through several examples the application of the workflow to biotechnological relevant problems, including the identification of alternative biosynthetic routes through enzyme promiscuity; or the development of biosensors. We demonstrate in that way the ability of the workflow to streamline retrosynthesis pathway design and its major role in reshaping the design, build, test and learn pipeline by driving the process toward the objective of optimizing bioproduction. The RetroPath2.0 workflow is built using tools developed by the bioinformatics and cheminformatics community, because it is open source we anticipate community contributions will likely expand further the features of the workflow.

**Highlights:**

- State-of-the-art Computer-Aided Design retrosynthesis solutions lack open source and ease of use
- We propose RetroPath2.0 a modular and open-source workflow to perform retrosynthesis
- RetroPath2.0 computes reaction network between Source and Sink sets of compounds
- RetroPath2.0 is distributed as a KNIME workflow for desktop computers
- RetroPath2.0 is ready-for-use and distributed with reaction rules

**Funding:** This work was supported by the French National Research Agency [ANR-15-CE1-0008], the Biotechnology and Biological Sciences Research Council, Centre for synthetic biology of fine and speciality chemicals [BB/M017702/1]; Synthetic Biology Applications for Protective Materials [EP/N025504/1], and GIP Genopole.

## 1. Introduction

Despite the increasing number of small molecules that are bioproduced, the process is still costly and rather slow. For instance, the metabolic engineering of artemisinic acid is claimed to have taken more than 130 person-years and about 10 years to complete (Paddon et al., 2013; Jay D. Keasling, 2014). Among the challenges that industrial biotechnology is facing to deliver sustainable solutions are 1) the reduction of R&D costs and 2) the bioproduction of a wider palette of compounds. To address these challenges, computational/experimental strategies where alternative metabolic pathways are first designed and assessed before being built and tested have been proposed (see reviews (Medema et al., 2012; Copeland et al., 2012; Hadadi and Hatzimanikatis, 2015; Lee and Kim, 2015)). While some computationally-driven strategies are making use of known metabolic reactions albeit not necessarily in the same species (Rodrigo et al., 2008; Moriya et al., 2010) others allow to design pathways that encompass novel reactions not stored in metabolic databases, these latter tools are making use of retrosynthesis algorithms (Marchant et al., 2008; Moriya et al., 2010; Henry et al., 2010; Yim et al., 2011; Liu et al., 2014; Carbonell et al., 2014a; Campodonico et al., 2014; Hadadi et al., 2016a).

Retrosynthesis algorithms take as input a set of metabolites, for instance the metabolites of a growth medium or the metabolites of a chassis strain model, and the set of target compounds to bioproduce. Ideally the target compounds could be any molecules of the chemical space. The algorithms generate retrosynthesis networks linking the target compound(s) (the source) to the metabolites of the chassis strain (the sink) through reactions.

Such retrosynthesis networks should be further processed to map or extract information relevant for the biological application. For instance, some algorithms can be applied to enumerate pathways (Carbonell et al., 2012) and rank them based on several criteria including enzyme availability and performance, product and intermediate compound toxicities (Planson et al., 2012) or the theoretical yield of the desired compound (Campodonico et al., 2014; Carbonell et al., 2014b; Cho et al., 2010; Liu et al., 2014). Interestingly, retrosynthesis networks exploitation is not strictly limited to retrosynthesis. Applications have been proposed to predict biodegradation routes (Hou et al., 2004; Oh et al., 2007; Finley et al., 2009) identify unknown compounds from the underground metabolism (Jeffryes et al., 2015), predict the transitions of labelled atoms in metabolic networks (Arita, 2003; Hadadi et al., 2016b), and design biosensing circuits for compounds for which no direct biosensors are known (Delépine et al., 2016). The main difference of the aforementioned applications lies in the definition of source and sink compounds sets; the current paper focuses on retrosynthesis but our solutions still stand for other applications requiring network generation.

One issue users of retrosynthesis-based solutions are facing is that algorithms and underlying data have not been fully documented and released. In most cases, authors provided fine-tuned webservers (Campodonico et al., 2014; Carbonell et al., 2014b; Jeffryes et al., 2015; Hadadi et al., 2016a) often filled with pre-generated data that focuses on some exemplar cases. Based on this information, it is difficult for users to grasp methods’ limitations, to improve them, or to exploit them for different uses. At a time when open-data principles gain more and more traction (Schofield et al., 2009; McNutt et al., 2016; Haug et al., 2017) we believe this lack of flexibility should be overcome.

In this spirit, we developed the RetroPath2.0 workflow on the KNIME analytics platform (Berthold et al., 2008) to answer the need for a modular and easy-to-use tool to predict reaction networks. Workflows have several advantages over scripting languages. A graphical user interface allows for rapid test and prototyping, even for users with little to no knowledge in programing. For instance, parallelization of tasks is inferred from workflow topology and does not need any special library or technical knowledge from the user. Once configured, workflows are readily deployable on all platforms where KNIME can be installed. KNIME workflows are popular in cheminformatics to prepare and analyse data, as shown by the number of extensions maintained by users in this field (Berthold et al., 2008; Warr, 2012). Thus, metabolic engineers beneficiate from a large panel of tools to analyse the chemical diversity and features of their data. As a matter of facts, RetroPath2.0 was developed using only community tools. We foresee it will make the workflow easier to modify and at the very least a good proof of concept of what can be done with workflows.

The current paper provides for the first time a simple workflow encompassing the main steps of the retrosynthesis process. We hereby review the main steps of retrosynthesis algorithms in order to demystify their use and shred lights on the shortcomings of current tools (Marchant et al., 2008; Moriya et al., 2010; Henry et al., 2010; Yim et al., 2011; Liu et al., 2014; Carbonell et al., 2014a; Campodonico et al., 2014; Hadadi et al., 2016a). We then outline our proposed solution through several applications in metabolic engineering and biosensor engineering. RetroPath2.0 is available in supplementary along a set of reaction rules and some classic metabolic engineering examples to test RetroPath2.0 features.

## 2. Theoretical background

### 2.1 Encoding reactions as reaction rules

The first challenge that retrosynthesis algorithms have to address is linked to the way reactions are encoded. Most retrosynthesis algorithms are based on reaction rules, but other strategies exist to encode reactions (Kayala et al., 2011; Latino and Aires-de-Sousa, 2011). A reaction rule generally depicts the change in bonding patterns when transforming a set of substrates (reactants) into a set of products. For retrosynthesis applications, rules are reversed such that one computes the substrates from the products.

Several solutions have been proposed to code for reaction rules, namely Bond-Electron (BE) matrices (Dugundji and Ugi, 1973), reaction SMARTS (Daylight, 2017), RDM patterns (Oh et al., 2007), and reaction signatures (Carbonell et al., 2013). Examples of coding systems are illustrated in Figure 1. We highlight below some key concepts to understand reaction rule encoding in a retrosynthesis context.

**Figure 1.**
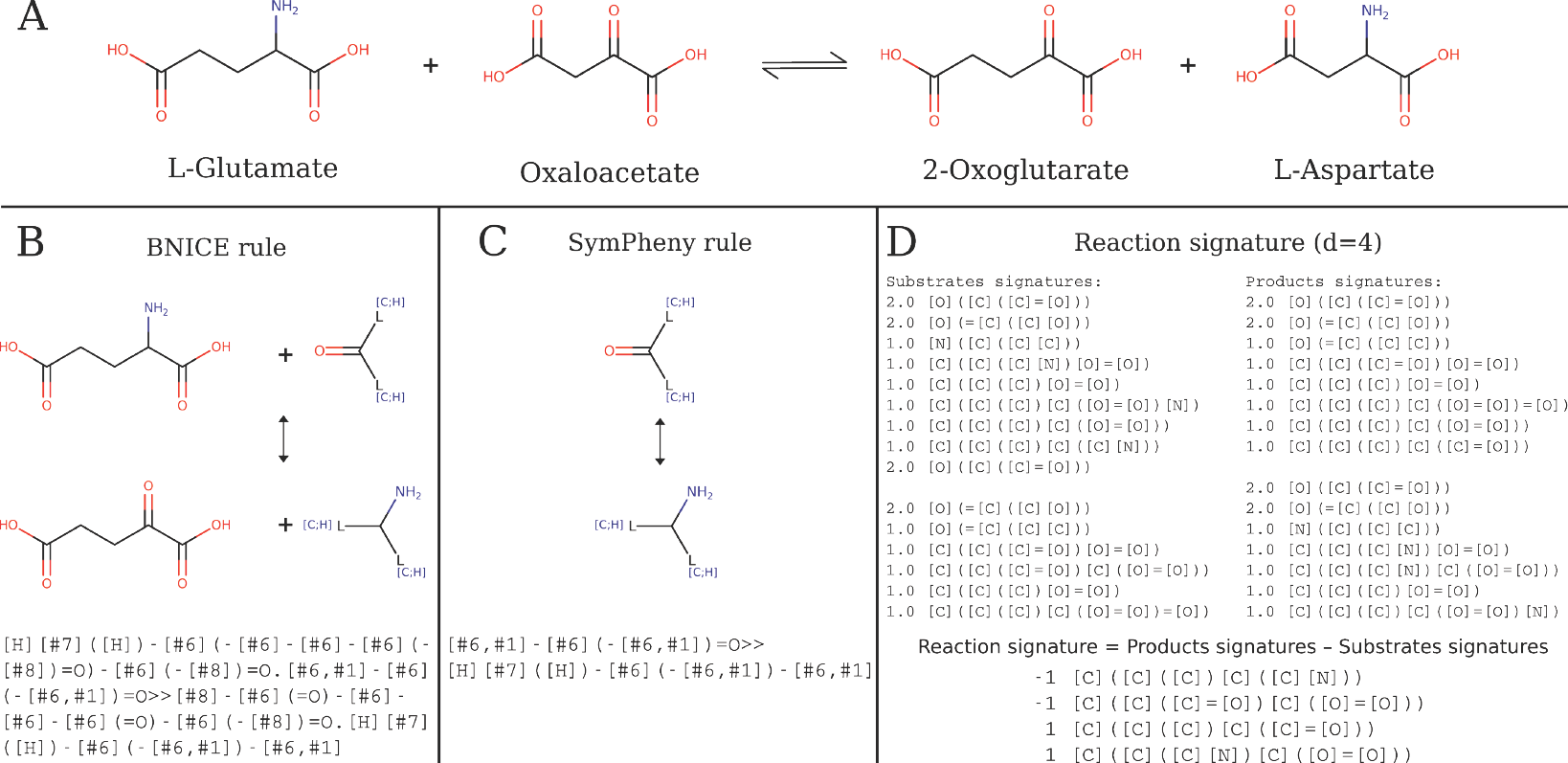
Example of reaction rules. A. Generalised reaction rules for the transaminase 2.6.1.1. B and C. BNICE rules and SimPheny rules were extracted from (Henry et al., 2010) and (Yim et al., 2011). These are the only rules with EC number 2.6.1. In both cases the rules are represented by SMARTS strings. D. The reaction signature rule was computed using the MolSig package, d represents the signature diameter. See (Carbonell et al., 2013) for definition and examples of signatures.

#### 2.1.1 Enzymatic promiscuity

Reactions for retrosynthesis applications should be modelled with a controlled degree of generalization for their substrates and products. Indeed, reaction rules containing a full description of substrates and products chemical structures cannot be applied on new compounds. This is the case for classic metabolic models and database and their lack of generalization prohibits the generation of novel pathways. The use of generalized chemical transformations is required in order to be able to predict new metabolic transformations. Such predictions are necessary since reaction databases are not complete (Altman et al., 2013; Chang et al., 2015) and side enzymatic activities are often underestimated.

This lack of knowledge on alternative enzymatic activities is currently a critical limiting factor for metabolic engineering since it has been estimated that 37% of *E.coli* K12 enzymes have a promiscuous activity for other substrates structurally similar to their main known substrate (Nam et al., 2012). In order to be able to generate new metabolic transformations (and new compounds) one thus needs to use generalized reactions to model enzymatic promiscuity, i.e. rules that can be applied to different substrates, and eventually on compounds absent from the databases. For instance, BNICE (Henry et al., 2010; Jeffryes et al., 2015; Hadadi et al., 2016a) and SimPheny (Yim et al., 2011) use a collection of reaction rules that, as depicted in Figure 1, can be applied to any ketones (including oxaloacetate) since their encoding is focused on the reaction centre.

#### 2.1.2 Identification of the reaction centre

The simplest way of controlling a degree of abstraction for reaction substrates is to encode the reactions around its centre. This requires compiling the list of atoms that belong to the reaction centre, i.e. atoms that change their configuration when the reaction is applied (panel B in Figure 2). Atoms changing configuration are those attached to bonds that are broken, formed, or are changing order, as well atoms for which charge and stereochemistry is changing when the reaction is taking place.

**Figure 2.**
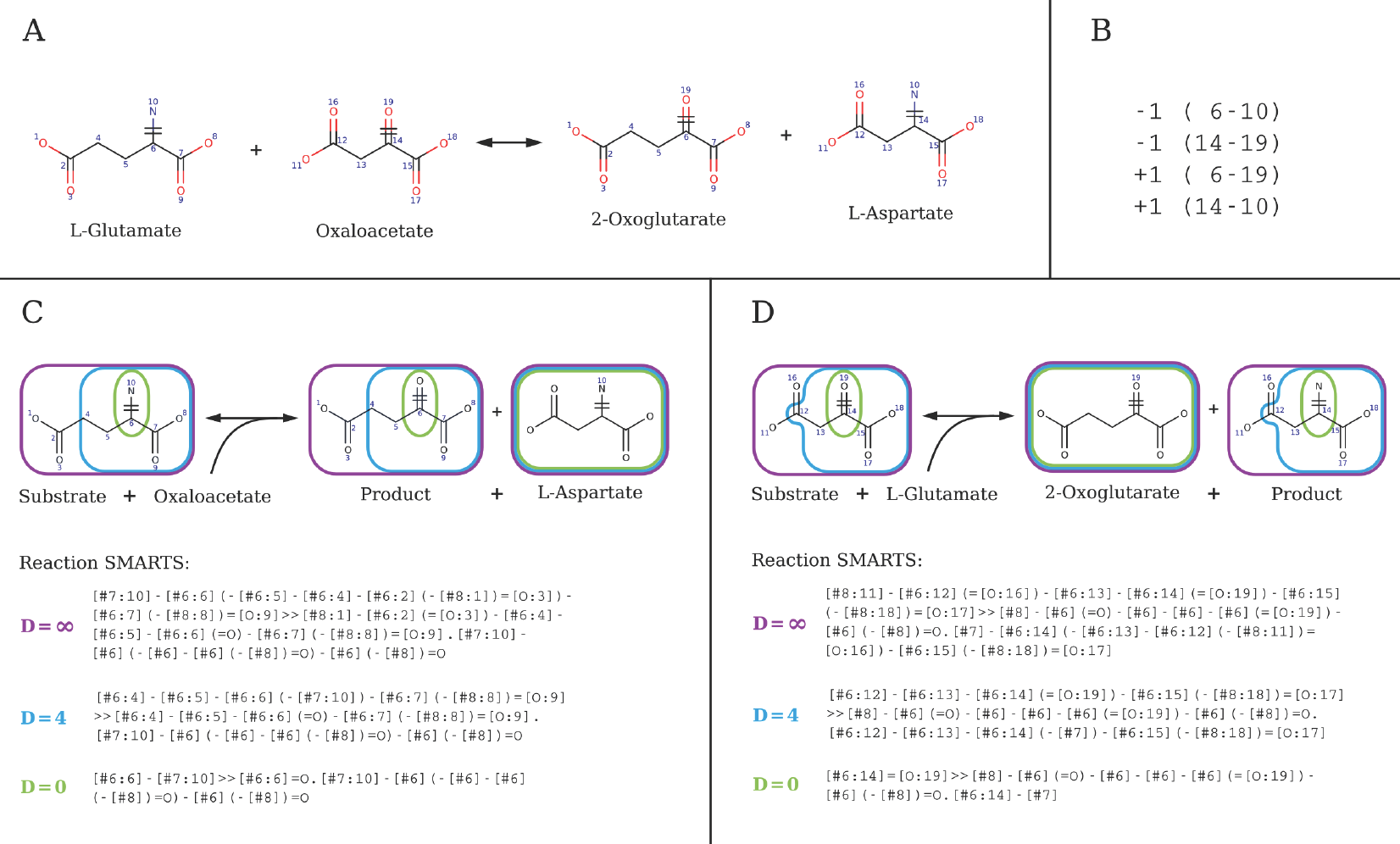
RetroPath2.0 rules and corresponding SMARTS for reaction 2.6.1.1 at various diameters. A. Full reaction 2.6.1.1 with atom mapping. B. The list of broken bonds (-1) and bonds formed (+1) is given by their atom numbers. C. The corresponding SMARTS for the component modelling promiscuity on L-glutamate: Substrate + Oxaloacetate = Product + 2- Oxoglutarate. D. The corresponding SMARTS for the component modelling promiscuity on oxaloacetate: Substrate + L-Glutamate = L-Aspartate + Product. C and D. Rules are encoded as reaction SMARTS and characterized by their diameter (∞ purple, 4 blue or 0 green), that is the number of bonds around the reaction centre (atoms 6, 10 and 14, 19) defining the atoms kept in the rule. This allow for a controlled and flexible modelling of enzymatic promiscuity. Note that for the case of 2.6.1.1 the co-product is always the same (C: L-aspartate; D: 2- oxoglutarate) but that is not always the case, depending on the connectivity of the atoms belonging to the reaction centre.

Reaction rules used in retrosynthesis generally require a solved Atom-Atom Mapping (AAM, see panel A in Figure 2) between the atoms of the substrates and those of the products to identify the reaction centre of the reaction (Hou et al., 2003; Hatzimanikatis et al., 2005; Oh et al., 2007; Cho et al., 2010; Liu et al., 2014). The AAM problem is equivalent to the Maximum Common Substructure, or the subgraph isomorphism problem which turns out to be NP-hard (Chen et al., 2013). Avoiding the use of AAM to generate rules is nevertheless possible in some cases, as it was originally shown by a previous version of the RetroPath algorithm based on fingerprint subtraction (Carbonell et al., 2014a) (see Figure 1).

Importantly, if encoding the reacting centre is necessary, it may not be sufficient to properly define a reaction catalysed by an enzyme since other atoms far from the reacting centre could be involved in the ligand binding as well. To palliate this problem, the definition of the reacting centre is extended to neighbour atoms, either systematically at a predefined bond-distance (diameter, panel C and D Figure 2) or based on expert-knowledge.

#### 2.1.3 Systematic rule generation

Reaction rules can be in principle computed by processing the set of reactions stored in metabolic databases. However there are some difficulties associated with this task and exhaustive rules generation is certainly another challenge for retrosynthesis. We can distinguish two main philosophies to systematically encode enzymatic reactions.

The first approach consists in encoding a small set of generalist rules guaranteed by a model to cover all possible class of reactions. According to the Enzyme Commission (EC) nomenclature all reactions that belong to the same third level EC number should follow the same chemistry, while the fourth and last level is for disambiguation (International Union of Biochemistry and Molecular Biology Nomenclature Committee and Webb, E.C, 1992). Both SimPheny and BNICE use the third EC number level to guide their reaction encoding effort. SimPheny (Yim et al., 2011) has 50 manually curated reaction rules, and the number of rules of BNICE systems are of the same order; 86 for (Henry et al., 2010), 198 for (Jeffryes et al., 2015), 722 for (Hadadi et al., 2016a). This approach is well-suited for manual curation, since the number of reactions to annotate is rather small but supposedly exhaustive in term of involved chemistry. Nonetheless, relying on EC numbers often requires to add exceptions since some reactions at a third level EC numbers do not share any common substructures and thus cannot be expressed by the same rule. For instance, the carbon-halide lyases class (EC 4.5.1.*) is composed of five fourth level reactions which all remove a chlorine atom, but some reactions also remove a primary amine from a substrate and replace it either by a double bonded carbon, a hydrogen, an oxygen atom or a more complex functional group (Supplementary Figure S1). Their number of substrates and products also varies. Clearly, these reactions cannot be encoded using a single BE matrix, a reaction signature, or an intelligible reaction SMARTS. Another need for exceptions arise from the fact that many reactions have no EC number assigned by the Commission (International Union of Biochemistry and Molecular Biology Nomenclature Committee and Webb, E.C, 1992).

The second approach, which is more data-driven, is to automatically compute rules for all available metabolic reactions by selecting only the atoms belonging to a sphere of fixed diameter around the reaction centre. This is the approach adopted by RetroPath (Carbonell et al., 2014a) and the workflow proposed in this paper. Using the procedure outlined in the caption of Figure 2, when applied to the MetaNetX database (Moretti et al., 2016) the number of rules returned is between 6,900 and 19,000 depending on the parameters used to model enzymatic promiscuity (diameter) for the 31,527 reactions stored in MetaNetX (MNXR identifiers, v.2.0). Interestingly, not only multiple generated rules can belong to the same EC class, but also a same rule can correspond to several EC classes. For instance, at diameter 4, three EC numbers (2.6.1.1, 2.6.1.17, 2.6.1.67) from three distinct reactions (resp. MNXR32641, MNXR32641, MNXR31792) are associated to the same rule depicted in Figure 1D (promiscuity on oxaloacetate, MNXM42).

#### 2.1.4 Cosubstrates, cofactors and coproducts

Another challenge for retrosynthesis algorithms is the need to handle reactions processing multiple substrates and/or multiple products. Dealing with multi-substrate reactions requires more computational resources in order to model enzymatic promiscuity for each combination of promiscuous substrates (Figure 2).

For these purposes, cosubstrates and coproducts that are currency cofactors (such as water, CO_2_, ATP, NADP, etc.) can be ignored from the rules under the assumptions that they are available in the cell and that there is no gain for retrosynthesis analysis in modelling promiscuity on them.

Nonetheless, there are still many metabolic reactions that are multimolecular, even after removing currency cofactors. MetaNetX version 2.0 (110,000 compounds after canonicalization and 31,527 reactions) comprise 42% of reactions that remains multimolecular after removing currency cofactors. Metabolic databases such as MetaCyc or KEGG do identify main substrates and products, albeit not in all cases, and 29% of reactions in MetaCyc have multiple main substrates and 27% have multiple main product and 15% have both. A good example of such reactions are transaminases (EC class 2.6.1). There are 178 reactions in that class in MetaCyc, most of them involve two substrates and two products. Cofactors in 2.6.1 reactions are glutamate and oxoglutarate in 51% of the cases, but other cofactors are found such as oxaloacetate, 2-oxobutanoate, oxoglutaramate, oxooctonal, glyoxylate, pyruvate, glutamine, oxosuccinamate, or butamine. Clearly reactions of class 2.6.1 admit multiple substrates and products and they all vary from one reaction to another, thus these reactions cannot all be coded as monomolecular transformations (R_1_C(=O)R_2_→R_1_C(NH_2_)R_2_) in the way that is done in (Yim et al., 2011) nor they can be coded in the form Glutamate + R_1_C(=O)R_2_ → Oxoglutarate + R_1_C(NH_2_)R_2_, (where R_1_ and R_2_ can be C or H) as done in (Henry et al., 2010).

Another issue that retrosynthesis algorithms need to overcome is to handle multiple products when the reactions are reversed (as they should be in any retrosynthesis process). Indeed when a reaction proceeds forward one assumes that the substrates are readily available and this is generally the case when moving down through a metabolic pathway, where the substrates of any given reactions are the products of upstream reactions. In retrosynthesis, the products become the substrates of the reversed reaction, and these substrates are not necessarily known. To illustrate this issue let us consider the reversed reaction T + P ⇒ S, where T is our retrosynthesis target. T is known but not P, in principle we should apply the rule to any compound P of the chemical universe (since P is not necessary a known metabolite). This solution is of course not practical. To palliate this issue, RetroPath do not reverse multiproduct reactions but construct an extended metabolic space using reaction rules fired on the metabolites chassis strains. The other retrosynthesis algorithms do not explicitly address this issue, albeit mono product reactions can certainly be reversed, and in principle, all SimPheny rules can be used for retrosynthesis purposes.

As summarized in Table 1 below, SimPheny (Yim et al., 2011) does not deal with multiple substrates and products as all rules are monomolecular, BNICE in (Henry et al., 2010) handles partially the problem, as 70 out of 86 reactions are multimolecular but only 4 reactions have multiple substrates different than cofactors (compared to 35% in MetaNetX). GEM-Path and RetroPath work with rules handing multiple substrates and multiple products, thus reflecting better the complexity of metabolic reactions. Nonetheless, RetroPath allows only one substrate at a time to undergo promiscuity modelling so that reaction prediction remains tractable.

**Table 1.** Retrosynthesis networks generation tools.

|  | Reaction rules calculation | Rules coverage | Number of rules | Reaction rule specificity | Multiple product & substrates | Enzyme sequence search | Combinatorial complexity | Availability |
| --- | --- | --- | --- | --- | --- | --- | --- | --- |
| SimPheny (BioPathway Predictor) (Yim et al., 2011) | Computed from 3rd EC level followed by manual curation | All metabolic reactions | 50 | Fixed | No | No | Controlled by network and molecules size | No |
| BNICE (Hadadi et al., 2016a) | Automated from KEGG followed by manual curation | All metabolic reactions | 722 | Fixed | Yes | No | Controlled by network size | Web server |
| PathPred (Moriya et al., 2010) | Automated from the KEGG RPAIR database | Xenobiotic degradation and biosynthesis of secondary metabolites | 853 (degradation)<br>1126 (biosynthesis) | Fixed | No | No | Controlled by similarity | Web server |
| GEM-Path (Campodonico et al., 2014) | Computed from 3rd EC level | All metabolic reactions | 443 | Fixed | Yes | No | Controlled by similarity and thermodynamics | No |
| METEOR (Marchant et al., 2008) | Knowledge-based expert system (Lhasa Ltd.) | All metabolic reactions? | 357 | Fixed | No? | No | Controlled by “reasoning” rules | Commercial |
| Reverse Pathway Engineering (THERESA) (Liu et al., 2014) | Automated from the BioPath database (Molecular Networks GmbH) | All metabolic reactions | 3,516 reference reactions | Fixed | Yes | Comparative genomics | Manual compound selection | Commercial |
| RetroPath (Carbonell et al., 2014b) | Automated computation on MetaCyc | All metabolic reactions | between 3,000 (d=4) and 5,000 (d=14) | Variable, controlled by diameter | Yes | Post-process using machine learning | Controlled by diameter | Web server |
| RetroPath2.0 (this study) | Automated computation on MetaNetX | All metabolic reactions | between 6,900 (d=2) and 19,000 (d=16) | Variable, controlled by diameter | Yes | Embedded using sequence clustering | Controlled by diameter and enzyme score | Open source |

### 2.2 Building (retrosynthesis) reaction network

In all algorithms listed in Table 1, retrosynthesis maps are constructed by applying reaction rules in an iterative fashion starting from a source set of compounds until the molecules in a sink set of compounds are found in the map. In the context of metabolic engineering, if the rules are applied in a forward manner, the source set is composed of the native metabolites of the chassis strain and the sink set are the molecule we wish to produce. If the rule are applied in a reverse manner then the source set are the molecules to be produced and the sink set are the metabolites of the chassis. One bottleneck that all algorithms face is computation complexity due to the combinatorial explosion of the number of reactions predicted from the rules. This is true disregarding if the reactions are applied in a forward or reverse manner. As an example, let us assume we wish to perform retrosynthesis for some FDA approved drugs in *E. coli.* In the reaction list we have at our disposal there is one for reversed hydro-lyases (i.e. reversed 4.2.1). According to (Henry et al., 2010) the rule for that reversed reaction is R1C(=O)C(R2)=C(R3)R4 + O-R5 → R1C(=O)C(R2)-C(R3)(R4)OR5, where R5 can be C, H, O, and S and all other Rs can be any atoms. Assuming R1C(=O)C(R2)=C(R3)R4 is the main substrate (our drug target) and O-R5 the cosubstrate, 68 FDA approved drugs from DrugBank contain the first substructure. If we restrict the cosubstrate to be in the *E. coli* model iJO1366 then 653 metabolites out of 810 compounds in the model contain the second substructure, while 50,810 compounds from MetaNetX will pass the substructure test. Taking Vitamin C as an example of a DrugBank compound that passes the substructure filter, one finds 1,883 unique products when applying the reversed rule 4.2.1 to Vitamin C and *E. coli* metabolites and 343,177 products when the cosubstrate is in MetaNetX. There are more products than substrates because for some substrates the reversed rule 4.2.1 applies to more than one location.

As already mentioned, for a given retrosynthesis target one needs to apply all rules to the target, all rules to the products obtained by application of the reversed reactions to the target, and so on until a predefined stop condition occurs (often the number of iterations). Clearly, if reaction rules generate more than 1,000 products even with 50 rules the problem starts to be challenging -if not impossible- to manage computationally after 2 or 3 iteration steps.

Strategies are needed in order to cope with that complexity. RetroPath proposes a solution where reactions are scored according to their ability to retrieve enzyme sequences catalysing substrate to product transformations. Reactions below a predefined score are removed from the retrosynthesis map. For any given reaction the sequence scores are computed by machine learning using a technique named tensor product. The machine learning tensor product is trained on all known pair enzyme sequence × (substrate, product) using Support Vector Machines (Faulon et al., 2008) or Gaussian Processes (Mellor et al., 2016). GEM-Path (Campodonico et al., 2014) proposes another strategy where for each reaction the substrates are accepted if they are similar enough to the substrates of the reference reactions.

We detail in the next sections a new implementation of RetroPath to predict reaction networks and perform retrosynthesis among other applications. RetroPath2.0 address the challenges listed above with a special attention to remain easy to use and modify by end users, unlike tools developed so far. In that sense, both the encoding of reactions into generalized rules and the actual use of those rules to predict new reactions depend strictly on resources developed by the community.

## 3. Method

The workflow proposed in this section can a priori be used to all systems presented in Table 1 to construct retrosynthesis maps as long as reaction rules can be coded by reaction SMARTS. As examples, we provide such set of rules extracted from the SimPheny, BNICE and RetroPath systems.

Our computational methods make use of in-house algorithms (Carbonell et al., 2014a), RDKit routines (Greg Landrum, 2016) and KNIME nodes (Berthold et al., 2008). They have been implemented in the form of a KNIME workflow -called RetroPath2.0- (Figure 3) that we provide in the supplementary materials, in addition of sets of rules, examples, and useful data files.

**Figure 3.**
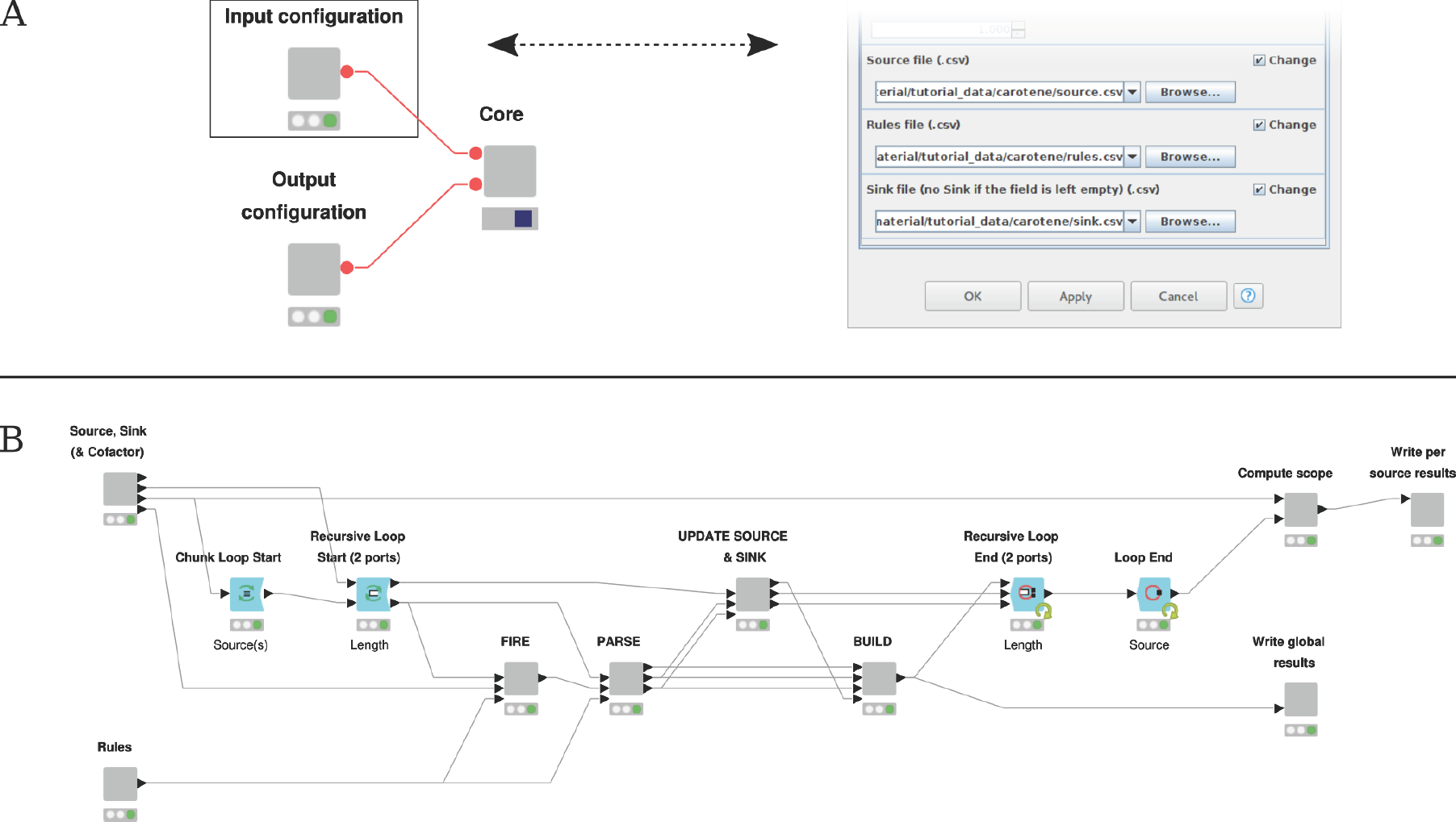
RetroPath2.0 KNIME workflow. A. Main panel view (left) and input configuration window (right) that allow the user to set up parameters. B. Inner view of the “Core” node where the computation takes place. The “Source, Sink…” and “Rules” nodes parse the source, sink and rules input files provided by the user and sanitize data so that it can be processed by downstream nodes. The outer loop (“Source” loop) iterates over each source compounds, while the inner loop (“Length” loop) allows to iterate the process up to a maximum number of steps predefined by the user. The nodes (i) “FIRE”, (ii) “PARSE”, (iii) “UPDATE SOURCE…” and (iv) “BUILD” are sequentially executed at each inner iteration. Respectively, they (i) apply all the rules on source compounds, (ii) parse and sanitize new products, (iii) update the lists of source and sink compounds for the next iteration and (iv) merge results that will be written by the node “Write global results”. Once the maximum number of steps is reached (or no new product is found), the “Compute scope” node identify the scope linking each source to the sink compounds, then these results are written by the node “Write per source results”. Only the main nodes involved in the process are shown.

### 3.1 Reaction rules

RetroPath2.0 uses reaction SMARTS to encode reactions. It is a SMIRK-like reaction rules (Daylight, 2017) format defined by RDKit and is mostly compatible with other tools (see Figure 1).

#### 3.1.1 Collected reaction rules

Rules for SimPheny and BNICE were extracted from (Yim et al., 2011) and (Henry et al., 2010) respectively, and manually entered by using Chemaxon Marvin Sketch software products (15.5.18, 2015, http://www.chemaxon.com). For each rule atom mapping was calculated by Marvin Sketch and the resulting rules were stored in SMARTS format as in Figure 2.

#### 3.1.2 Generated reaction rules

We used MetaNetX version 2.0 (Moretti et al., 2016) as the reference database for metabolic reactions that we encoded in rules. MetaNetX is a meta-database that compiles into a single reference namespace both reactions and metabolites extracted from main metabolic databases such as KEGG, Metacyc, Rhea or Reactome.

Reactions can contain many substrates and many products. We performed an Atom-Atom Mapping (AAM) using the tool developed by (Rahman et al., 2016) on all MetaNetX reactions (Figure 2A). We filtered out transports reactions and those involving compounds with incomplete structures (class of compounds, R-groups, etc.). Stereochemistry was removed.

Multiple substrates reactions were decomposed into components (panel C and D in Figure 2). There are as many components as there are substrates and each component gives the transformation between one substrate and the products. Each product must contain at least one atom from the substrate according to the AAM. This strategy enforces that only one substrate can differ at a time from the substrates of the reference reaction when applying the rule (see section 2.1.1 enzymatic promiscuity modelling).

Next step consisted in computing reactions rules as reaction SMARTS for each component. We did it for diameters 2 to 16 around the reaction centre (panels C and D in Figure 2) by removing from the reaction components all atoms that were not in the spheres around the reaction centre atoms.

We extracted more than 24,000 reaction components from MetaNetX reactions, each one of those leading to a rule at each diameter (from 2 to 16).

We provide in Supplementary (at https://www.myexperiment.org/workflows/4987.html) a subset of 14,300 rules for *E.coli* metabolism, both in direct and reversed direction. The rules were selected based on the MetaNetX binding to external databases and the iJO1366 whole-cell *E. coli* metabolic model (Orth et al., 2011).

### 3.2. Building (retrosynthesis) reaction networks between two pools of compounds using the RetroPath2.0 workflow

The RetroPath2.0 workflow essentially follows an algorithm proposed by some of us (Carbonell et al., 2011). After removing all source compounds already in the sink set, the workflow applies the rules to each of the compounds of the source set. For each compound, the products are computed using the RDKit KNIME nodes (Greg Landrum, 2016). Products are standardised and duplicates are merged. All pairs substrate-product are added to the growing network along with the reaction rules linking them.

In the next iteration, the set of products become the new source set. However, before iterating, the workflow removes from the new source set all compounds that belong to the sink (as these are already solutions and there is no need to iterate) and the workflow adds the product set to the sink in order to avoid applying reactions on the same products during subsequent iterations. Consequently, the workflow computes only the minimal routes between source and sink, i.e. routes in which all reactions are essential for their viability, and thus minimizes the number of enzymes to be added to a chassis strain when implementing the pathways. This feature can be ignored by not specifying a sink for the first iteration.

The RetroPath2.0 workflow iterates until a predefined number of iterations is reached or until the source set is empty. The final produced graph is composed of the list of links between substrates and products annotated with their corresponding reaction rule. Products belonging to the sink are annotated as such.

Note that the iterative process can reveal itself to be quite computationally demanding. To tackle this issue, RetroPath2.0 has a feature to bias the reaction space exploration toward compounds generated by trusted rules, using a rule-wise penalty score. If too many compounds are generated to be handled at once, only a predefined number of compounds with the lowest penalties according to their generating rules are kept in the new-source of the following iteration. Of course, both the definition of the penalty and maximum number of compounds to keep are critical and fall within the responsibility of the user. As described next, the rules we provided are scored to optimize pathway *in vivo* feasibility by penalizing rules associated to enzymes with inconsistent sequence annotation.

### 3.3. Score rules by enzyme sequence consistency

Predicted reactions in the final graph generated by the RetroPath2.0 retrosynthesis workflow need to be associated with enzyme sequences in the final engineering of the pathways. The selection of such sequences should look for a trade-off between the specificity of the reaction rule and the information available in enzyme databases for the reaction through the EC classification. Whereas the EC classification has traditionally provided a hierarchical numerical classification of enzyme-catalysed reactions to progressively describe reactions in finer detail, RetroPath2.0 introduces a similar hierarchical classification that is controlled by the diameter used in rule generation. In some cases the diameter of the reaction rule found by the RetroPath2.0 workflow might be high, i.e. highly specific to that reaction. However, it often occurs that there is no annotated enzyme sequence for the rule. In order to find some candidate sequences, we look into reactions that are close according to the EC hierarchy for each EC class containing at least one instance of the rule at given diameter. Traversing both rules diameter hierarchy and the underlying EC classes allows the selection of plausible sequence candidates for each reaction rule.

We compiled the set of Uniprot sequence identifiers annotated for reactions by looking at the cross-link annotations in MetaNetX for Rhea and MetaCyc databases. In total 208,980 sequences from 5,388 organisms were associated to 7,793 reactions. At a given diameter of the rule, we iteratively assigned sequences to rules. First, reactions with annotated sequences were collected for each generated rule. Since a rule can represent one or more reactions at a given diameter, sequences coming from different reactions sharing the same rule were aggregated into a single set for that rule. These direct annotations only provided a partial coverage for the total rules in the database. For instance, at diameter *d* = 8, there were 7,898 orphan rules, i.e. rules that were generated from reactions lacking sequence annotation (Supplementary Table 1). Similarly, there were 6,280 orphan reactions at diameter *d* = 8. In order to increase the coverage, we considered the EC class of the reaction when such information was available. Sequences associated with reactions sharing strictly the same EC class were combined together. Adding together such annotations for the same EC class fixed issues related to partial annotations for the less common reactions. In that way, the number of orphan rules was significantly reduced to 1,719, which is approximately a 13% of the total rules. Similar ratios were observed for reactions.

For the orphan rules having no sequence annotation after considering the EC class of the reactions, we followed the strategy of reducing the specificity of the EC class by reducing the number of digits. In other words, if a rule had no annotation based on the EC class at 4 digits, we looked at reactions that shared same EC class at 3 digits with one reaction associated with the rule and so on until we found sequence annotations. Notably, a sharp decrease on the number of orphan rules already occurred at the level of three digits of the EC class. The remaining orphan rules, less than 1%, was eventually annotated once we reduced the specificity from 3 digits down to 1 digit in the EC class.

We should emphasize that in the procedure described below, sequence annotations that merged multiple EC classes sharing same initial digits were only used for those cases where no sequence information was available at higher EC class levels. This annotation from higher to lower specificity in the set of sequences associated with the rules depending on known sequences allowed us to score the rules. A rule that has associated sequences with low diversity should in general correspond to cases where the sequence information is highly specific to that rule. As the diversity of sequences increases the specificity of those sequences to their associated rules becomes lower. We evaluated such degree of specificity by considering the degree of clustering of the sequences associated with the rules. Clustering of the sequences was performed by using Cd-hit (Li and Godzik, 2006). According with this algorithm, our database of 208,980 amino-acid sequences was clustered into 22,221 clusters for a similarity threshold of 0.5. We used a penalty score for the rules based on the number of sequence clusters *n_rule_* contained in the sequences selected for a given rule:

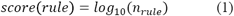

where the logarithm is applied for regularization. A penalty score of 0 implies high specificity, as this means that all sequences belong to a single cluster, while high penalty scores imply multiple clusters and therefore low specificity in the sequence annotation.

### 3.4. Enumerating pathways between two pools of compounds

The lists of pathways linking (i) a pool of source compounds to (ii) a pool of sink compounds are computed running an algorithm we developed earlier (Carbonell et al., 2014a). This algorithm consists of the following steps for a given source compound. (1) Compute the scope, a subset of predicted reactions between the sink compounds and the set of source compounds. The scope represents the set of compounds and reactions that are involved in at least one pathways. It is computed in a two steps search. First the forward step starting from source compounds finds all reachable compounds that are producible through reactions. Secondly the backward step starting from the sink compound adds to the scope all reactions that can be involved in at least one producible-pathway. (2) Build the stoichiometric matrix. The stoichiometric matrix describes the directed subnetwork involving the set of compounds and reactions identified at the scope step, starting from the source compounds. (3) Enumerate elementary flux modes. An elementary mode corresponds to a minimal unique set of reactions that (i) verified the stoichiometric constraints of the network and (ii) is able to carry non zero-fluxes at the system’s steady-state (Schuster et al., 2000). In order to efficiently compute elementary modes, stoichiometric matrix dimension is generally reduced through lossless compression. Only enumerated flux modes linking source compounds to the sink compound are kept in order to form the final list of pathways. These three steps are performed iteratively for each source compound.

RetroPath2.0 computes the scope for each queried compound. It can be visualized and explored to retrieve the pathways thanks to ScopeViewer, a humble web-application that we provide in Supplementary (at https://www.myexperiment.org/workflows/4987.html). Note that the provided workflow does not explicitly extract the pathways.

## 4. Results

We validated our set of rules with RetroPath2.0 by checking that they were able to reproduce known metabolic space, and that they could be used to perform reaction classification. The capability of RetroPath2.0 to perform retrosynthesis was confronted to *in vivo* experiments by counting the number of bioproduction pathways found for targets extracted from a database of metabolic engineering successes. We finally emphasized the versatile usage of RetroPath2.0 by an original application to design biosensors.

### 4.1. Rules validation

The quality of the output of the workflow depends largely on feeding it with the proper set of reaction rules. Some authors (Henry et al., 2010; Yim et al., 2011) have published sets of rules that already constitute an initial testbed. We collected those in addition of a set of SMARTS rules that we compiled for all reactions of the last *E. coli* whole-cell model (Orth et al., 2011) based on MetaNetX cross-references. All rules were checked to ensure they could be used with the workflow and yield at least one product.

#### 4.1.1. Coverage of known metabolic space

In order to check the potency of the rules, i.e. that they could indeed be used to predict reactions, we tried to retrieve all reference reactions of MetaNetX from the rules. We compared three dataset of monosubstrate rules according to their origin: SimPheny (Yim et al., 2011), BNICE (Henry et al., 2010) and RetroPath2.0. To make a fair comparison we selected from all MetaNetX reactions a subset of 13,000 reactions having an associated EC number and a structure for all its compounds (SimPheny and BNICE rules are based on EC numbers). We extracted from those 6,000 substrates and 7,000 products (MetaNetX identifiers) excluding cofactors. For each rule dataset, all rules were applied on the set of substrates using the workflow with default parameters. We counted the number of products that could be regenerated and the number of generated compounds that were referenced in MetaNetX.

Remarkably given the number of rules considered, 34% of MetaNetX products were recovered by SimPheny rules (50), and 41% by BNICE rules (86). They respectively generated 75,400 and 59,000 compounds, among which 5% and 7% could be found in MetaNetX and are thus connected to a biological database. Since RetroPath2.0 rules were generated from MetaNetX data we expected a better coverage over the products. This was indeed the case with 96% recovered products, and 37% of the 17,500 generated compounds found in MetaNetX.

The fact that RetroPath2.0 rules generates less compounds than the other tested sets of rules is explained by the differences in term of diameter used. RetroPath2.0 uses a flexible diameter, which by default ranges from 16 to 2, decreasing if no rule can be used on a substrate at higher diameters. This has for effect to prioritize more conservative results (higher diameter) while ensuring that broader promiscuity are tested in last resort (lower diameter). The few missed products originated from reactions that could not be encoded in rules due to atom-atom mapping issues.

Overall, product coverage shows us that RetroPath2.0 rules are able to reproduce most of MetaNetX products, hence most of what is known of the metabolic space.

#### 4.1.2. RetroPath2.0 rules for reaction classification

We evaluated the ability of our rules to perform automated reaction classification. To that end, reactions in the database that contained EC class annotations were grouped into their corresponding EC class at the third level. We then computed the similarity between reactions based on the signature content of their rules. For a given diameter *d,* each rule was decomposed into its elementary signatures (Carbonell et al., 2014a) and similarity between two given reactions *R_1_* and *R_2_* was computed by means of the Jaccard similarity coefficient *T^d^(R_1_,R_2_)* applied to the two reaction rules:

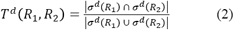

The previous expression ranges between 0 (minimum similarity) and 1 (maximum similarity) and has been often applied to compute similarity between compounds or even reaction that are described by binary fingerprints (EC-BLAST (Moriya et al., 2010; Campodonico et al., 2014; Rahman et al., 2014). The advantage and main difference of using rules with a selectable diameter is that we can compute the Jaccard similarity coefficient in function of the diameter *d*. That generates a sequence of monotonically decreasing similarities starting from 0 up to the maximum diameter of the reactants. Similarity of two reactions at diameter 0 contains the basic information about common patterns of bonds that were broken or formed in the two reactions. As we extend similarity to higher diameters, information becomes more specific to the substrates and products involved in each reaction.

In order to capture efficiently this feature of diameter dependence for Jaccard similarities between rules, we defined a global similarity parameter between reactions S(R_1_, R_2_) extended to a diameter range [0, *d*] as an exponentially increasing weighted sum of the Jaccard similarity coefficients:

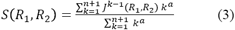

where *a* is a regularization parameter.

For each reaction in the database, we computed its corresponding rule and similarities based on a diameter range from 0 to 8. In total, rules were computed for 13,782 reactions contained in the database. We used *a* = 2 as regularization parameter.

We then tested the discriminant ability of using such reaction global similarity measure for reaction classification. Our tests were performed using the R package ROCR. We created a positive and negative set for each EC class. The positive set was formed by the set of reactions annotated for this EC class. A balanced training set was then built by randomly selecting from the negative set. For each EC class containing at least 10 data points, as well as for the total set of balanced training set we computed the area under the ROC curve (AUC), resulting in an overall AUC of 0.884 for diameter *d* = 8 (Fig. 4). Such performance values are slightly higher than the ones obtained by EC-BLAST (Rahman et al., 2014) by using fingerprint-based similarities, showing the ability of the rules as reaction classifiers.

**Figure 4.**
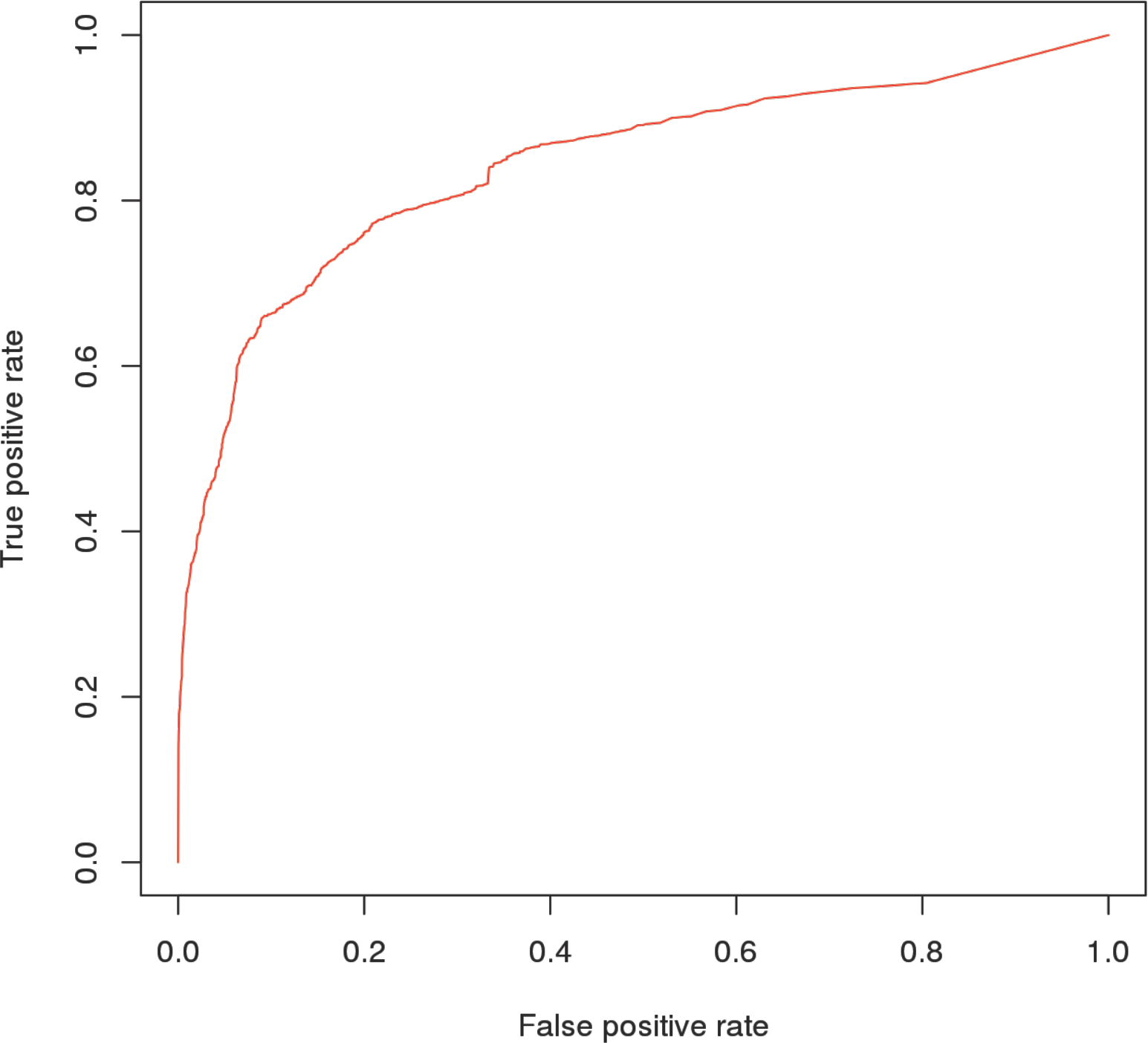
Receiving operating characteristic curves (ROC) curves for the rules of RetroPath2.0 of diameter *d* = 8.

#### 4.1.3. Score vs. specificity

The ability of substrate generalization of SMARTS rules can potentially be used to assess enzyme specificity. Enzyme specificity is an important factor that needs to be considered for metabolic pathway engineering. Moreover, several studies have shown that enzymes that can catalyse multiple reactions or can process multiple substrates have more evolvability capabilities than specific enzymes (Khersonsky and Tawfik, 2010; Nam et al., 2012; Orth and Palsson, 2012; Guzmán et al., 2015). Such property can be approached through our rules as they provide a means for representing chemical transformations for generalized substrates. The level of generalization of reactions and ultimately of their associated enzyme sequences could be therefore quantified using our rules. As described in Methods, one can define a specificity score by assessing the level of generalization of both the reactions and sequences having such reactions at a given rule diameter. The algorithm traverses both the reaction and sequence space in order to score reaction specificity and more specific rules get lower scores.

To evaluate the ability of the score to represent enzyme specificity, we have analysed a reference set of enzymes in *E. coli* that have been classified as either specific or generalist, i.e. if they can catalyse one or multiple reactions (Nam et al., 2012). For each gene, we took their associated reactions in the EcoCyc database (Keseler et al., 2013). Each reaction was mapped into their associated rule at several diameters *d.* The resulting scores for each gene were then aggregated. We mapped in total 787 *E. coli* genes, with 602 specific vs. 185 generalist enzymes, respectively.

Notably, the scores computed in that way, as shown in Figure 5, displayed the ability to differentiate between these two groups of enzymes, (t =−6.5144, p-value of 2.3e-10 for a Welch’s two sample t-test), with specific enzymes receiving lower ranking. We should note that the classification between specific vs. non-specific enzymes depends on the actual knowledge and degree of detail in the description of the reactions in the reference organism and therefore the list of generalist enzymes should be updated as long as new activities are discovered (Guzmán et al., 2015). For instance, we observed a clear outlier in the set of specific enzymes that received a high score based on rules and therefore we should expect wider specificity. This was the case of gene *phoA, b0383,* alkaline phosphatase EC 3.1.3.1. It turned out that this enzyme has been reported to have wide specificity (Yang and Metcalf, 2004) in agreement with the high score.

**Figure 5.**
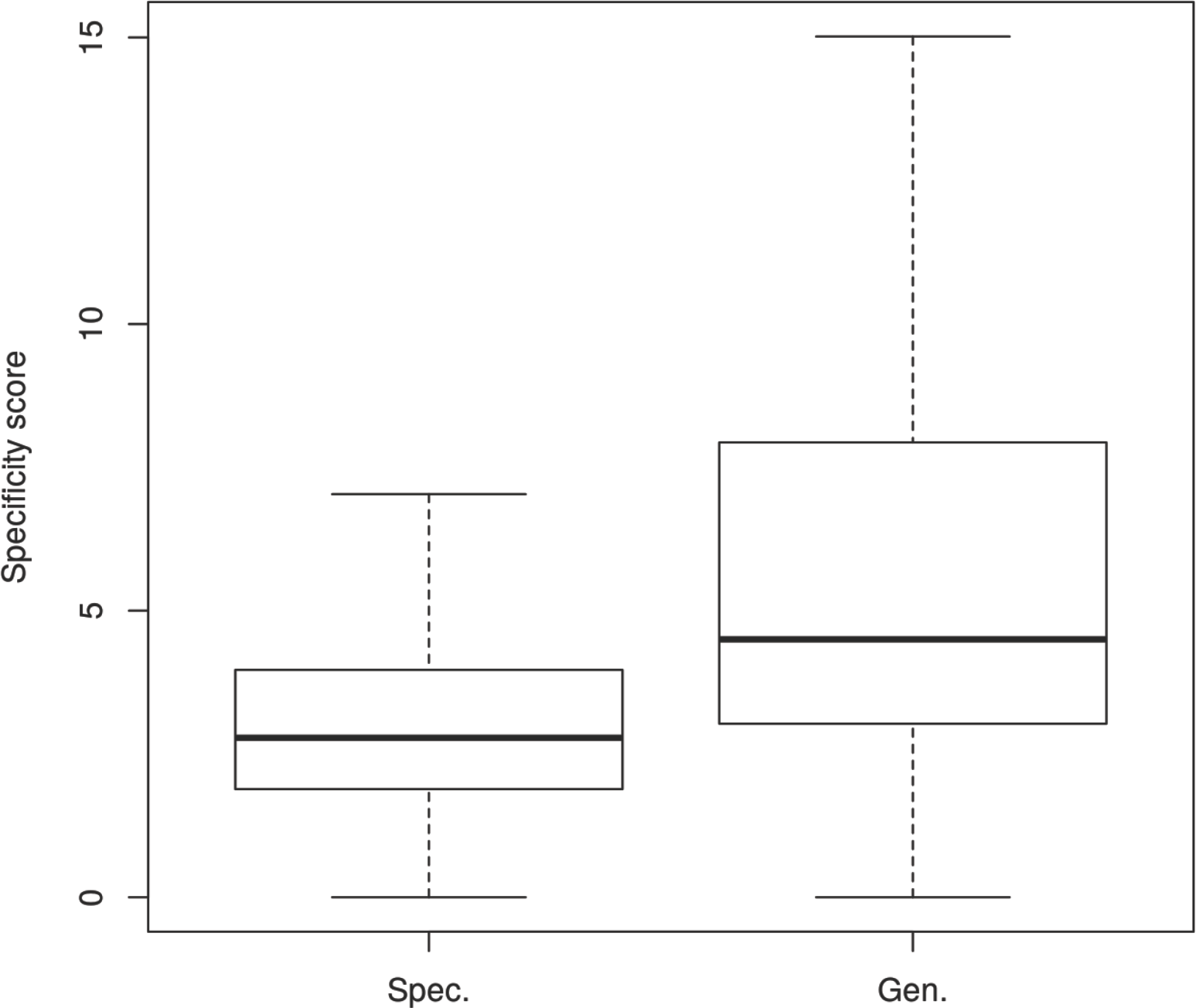
Box plot comparing the distribution of reaction scores for specialist and generalist enzymes in *E. coli.*

### 4.2 Workflow validation and applications

#### 4.2.1 Coverage of bioproduction pathways

The Learning Assisted Strain EngineeRing (LASER) database is a repository for metabolic engineering strain designs (Winkler et al., 2015). It stores more than 600 successful metabolic engineering designs (Winkler et al., 2016) that have been manually curated from the literature. Those examples are particularly appealing to test retrosynthesis features since they include an ideal dataset of authentic positive examples of bioproduction pathways, sometimes involving heterologous enzymes. We extracted all compounds targeted for production described in the LASER database (release f6ce080a8993) and used those to assess the ability of RetroPath2.0 to find retrosynthesis pathways for real-life applications when used with all the rules from MetaNetX.

The structures of the target compounds were inferred from their name by querying PubChem and ChemSpider. 160 compounds targeted for bioproduction were extracted from LASER. To complete further this dataset, we extracted 68 compounds (MBE dataset) published in Metabolic Engineering in 2016 (volumes 33 to 38), a period not covered by LASER. Both datasets have 203 distinct compounds once merged together based on their structure (standard InChI). Furthermore, we removed *E. coli* endogenous compounds that were used as our “sink”. Finally, 146 distinct compounds were collected to serve as “source” compounds.

Compounds from *E. coli* were extracted from iJO1366 whole-cell model (Orth et al., 2011) and MetaNetX cross-references. We ignored compounds that belong to so-called “blocked pathways” which are by definition impossible to produce or consume at steady-state in a metabolic model. Such compounds do not constitute a proper source (or sink) for retrosynthesis applications since a reaction explaining the compound availability in the chassis could be missing. We performed a flux variability analysis to identify them. Overall, we collected 962 MetaNetX identifiers of compounds belonging to *E. coli* that we provide in Supplementary along with their structure (InChI), at https://www.myexperiment.org/workflows/4987.html.

All results were generated with a maximum of five retrosynthesis iterations and a timeout of three hours by target on a recent desktop computer. Given those constraints, we successfully found at least one pathway for 81% of the targets (119/146), i.e. a set of reactions allowing the production of the target compound exclusively from *E. coli* endogenous metabolites. Interestingly, we found more than one pathway in most of the cases (104/119).

One of such compounds for which several pathways have been found is styrene. Styrene is a building block used in the fabrication of plastics (Isikgor and Becer, 2015). LASER references one pathway for the bioproduction of styrene from phenylalanine with heterologous enzymes in *E. coli* (McKenna and Nielsen, 2011; McKenna et al., 2015) and in *S.cerevisiae* (McKenna et al., 2014). RetroPath2.0 found this pathway (Figure 6, in red) along with five alternative one from *E. coli* endogenous compounds: 3-phenylpropionic acid, phenylacetaldehyde, and phenylpyruvic acid (Figure 6, resp. F, G, and H).

**Figure 6.**
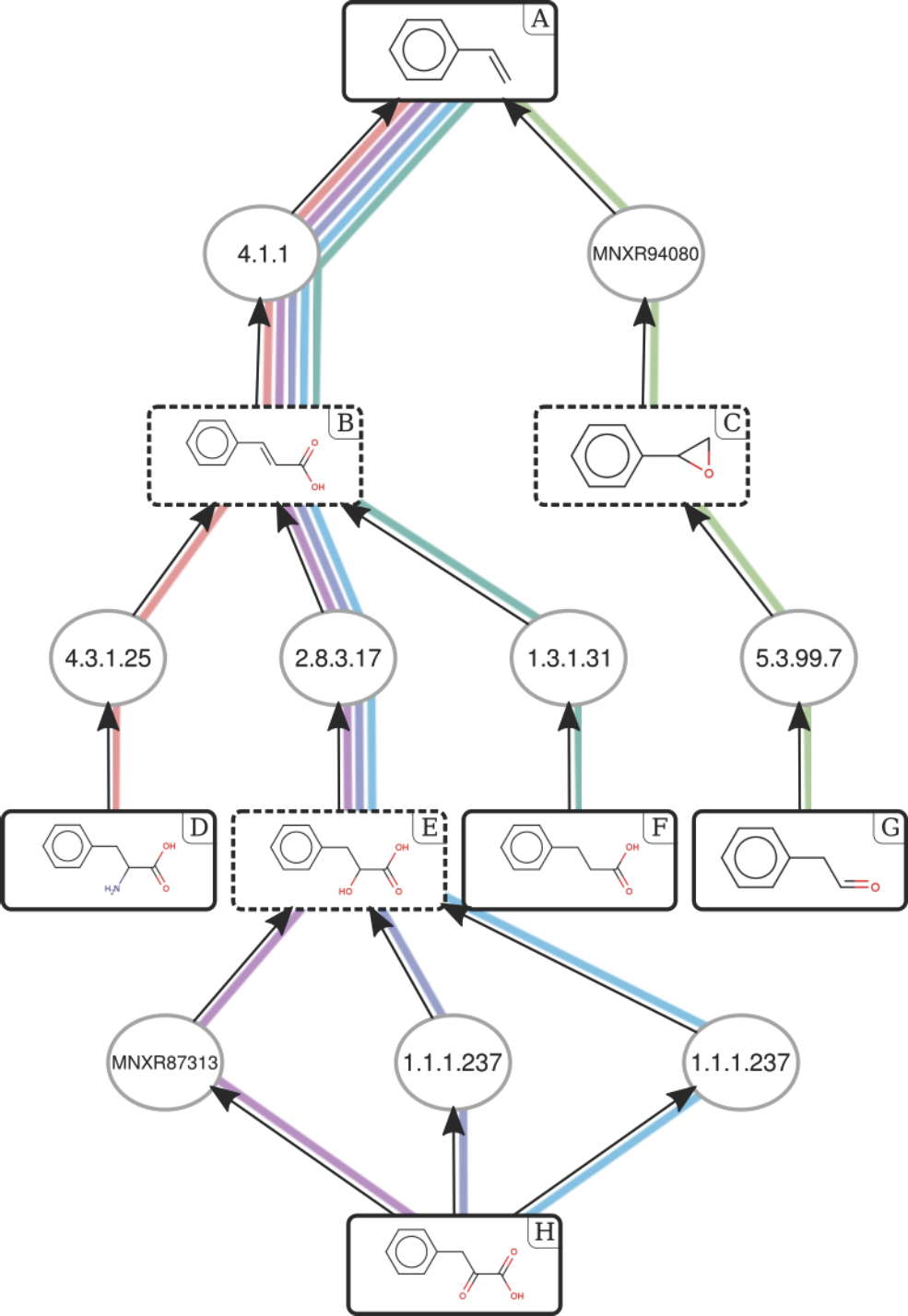
Enumerated pathways for the production of styrene. Each pathway is depicted by a distinct colour. Pathway referenced in (McKenna and Nielsen, 2011) is in red (D-B-A). Compounds are represented by their structures, and reactions by their EC numbers. Styrene and sink compounds are surrounded by a solid line, intermediates by a dashed line. A: styrene; B: phenylacrylic acid; C: styrene oxide; D: phenylalanine; E: 3-phenyllactic acid; F: 3-phenylpropionic acid; G: phenylacetaldehyde; H: phenylpyruvic acid. Cofactors have been removed for clarity; the whole scope is available in Supplementary at https://www.myexperiment.org/workflows/4987.html.

Another non-natural example for which several pathways are found is terephthalic acid (TPA). TPA is a non-natural commodity chemical widely-used for its ability to form synthetic fibres, and ultimately in the fabrication of polyesters such as PET. TPA is traditionally produced from p-xylene by synthetic chemistry processes (J. Sheehan, 2000). The p-xylene can eventually come from lignocellulosic biomass, making the TPA a bio-based compound in such cases (Isikgor and Becer, 2015). Interestingly, two enzymatic bioproduction pathways have been reported for TPA, and they follow the same chemical transformations as the ones from synthetic chemistry (J. Sheehan, 2000); one from p-xylene (Bramucci et al., 2001) in *Burkholderia* genus, and another from p-toluic acid in *Comamonas testosterone* (Wang et al., 2006). RetroPath2.0 retrieved those routes and proposed alternative shorter paths from endogenous *E. coli* compounds such as phenylalanin, phenylpyruvic acid, and 3-phenylpropionic acid (Fig. 7, resp. K, P, and M). To the best of our knowledge, those pathways have never been implemented *in vivo*.

**Figure 7.**
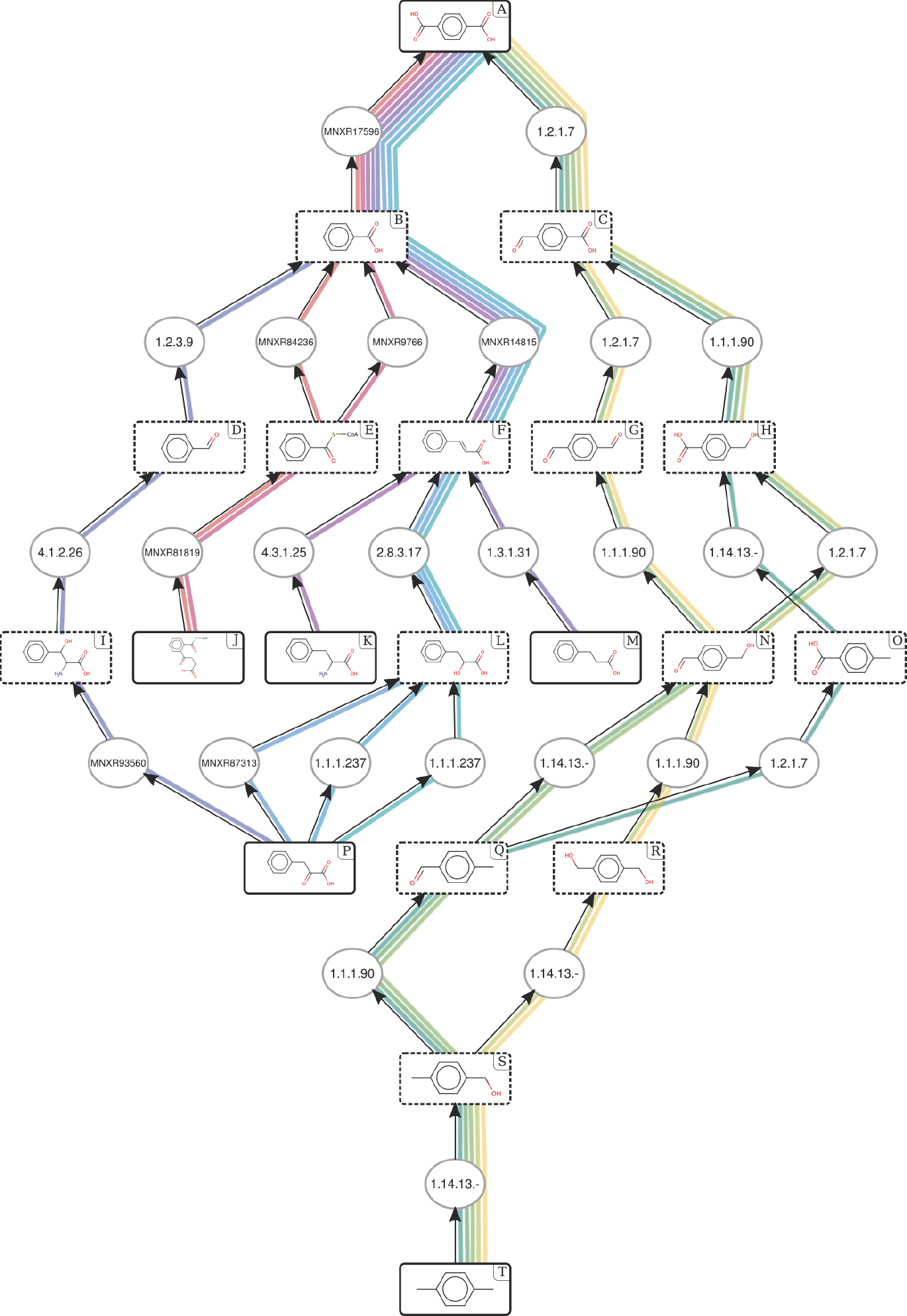
Enumerated pathways for the production of the non-natural compound terephthalic acid (TPA, compound A) from *E. coli*. Each pathway is depicted by a distinct colour. Pathway referenced in (Bramucci et al., 2001) is in teal blue (T-S-Q-O-H-C-A). Compounds are represented by their structures, reactions by their EC numbers. TPA and sink compounds are surrounded by a solid line, intermediates by a dashed line. Reactions with unknown EC number according to MetaNetX are referenced by their MetaNetX ID. A: terephthalic acid; B: benzoic acid; C: 4-formylbenzoic acid; D: benzaldehyde; E: benzoyl-CoA; F: phenylacrylic acid; G: terephthaldehyde; H: p-hydroxymethyl benzoic acid; I: 3-phenylserine; J: 2-succinylbenzoyl-CoA; K: phenylalanin; L: 3-phenyllactic acid; M: 3-phenylpropionic acid; N: 4-(hydroxymethyl)benzaldehyde; O: p-toluic acid; P: phenylpyruvic acid; Q: p-tolualdehyde; R: 1,4-benzenedimethanol; S: 4-methylbenzyl alcohol; T: p-xylene. Cofactors have been removed for clarity; the whole scope is available in Supplementary at https://www.myexperiment.org/workflows/4987.html.

Those results highlight the interest of RetroPath2.0 for retrosynthesis applications. RetroPath2.0 is able to reproduce validated pathways and propose new ones, both for natural and non-natural compounds. All results are provided in Supplementary (at https://www.myexperiment.org/workflows/4987.html).

#### 4.2.2 Detection of biomarkers through metabolic circuits

Asides from metabolic engineering, reaction network prediction algorithms can also be used to develop whole-cell biosensors. The typical synthetic biosensors (Khalil and Collins, 2010) currently being developed comprise a system capable of sensing a small molecule generally though allosteric interactions with RNA aptamers (e.g. riboswitches) or transcription factor (van der Meer and Belkin, 2010) and upon sensing express a reporter gene. In the context of medical diagnostic based on biomarkers detection, the main advantages of synthetic cell-based technologies over abiotic detection based on purified antibodies, nucleic acid hybridization, or metabolomics analysis are lower cost, improved stability, and the possibility to be ultimately used as a personal home healthcare device.

However, as of today, typical whole-cell biosensors are triggered by no more than half a dozen input signals. To palliate this shortcoming, we have recently proposed to expand the range of biologically detectable by systematically engineering sensing enabling metabolic pathways (SEMP) (Libis et al., 2016; Delépine et al., 2016), i.e., metabolic pathways that transform non-detectable chemicals into molecules for which sensors already exist. The SEMP method has been successfully benchmarked to engineer biosensors that detect pollutants, drugs and biomarkers such as benzoic acid and hippuric acid (Libis et al., 2016).

Here we further investigate the use of RetroPath2.0 to search all prostate cancer biomarkers that could potentially be detected using *E. coli* as a sensing device.

Prostate cancer biomarkers were retrieved from the Human Metabolome Database (HMDB) and scanned literature to select biomarkers in various physiological fluids: serum (Sreekumar et al., 2009; Zang et al., 2014; Li et al., 2016; Lima et al., 2016), urine (Sreekumar et al., 2009; Zhang et al., 2013; Struck-Lewicka et al., 2015; Fernández-Peralbo et al., 2016; Lima et al., 2016), and tissue (Sreekumar et al., 2009; McDunn et al., 2013; Lima et al., 2016; Huan et al., 2016). The above references gave a final list of about 800 small molecule biomarkers. Because biosensors are considered to be engineered in *E. coli* we removed all *E. coli* native metabolites, we also removed duplicates and biomarkers that could not be found in HMDB because of ambiguous names. The resulting sanitized set was composed of 421 biomarkers (provided in supplementary materials).

RetroPath2.0 was run taking as source all (non-*E.coli*) prostate cancer biomarkers, and as sink a list of 500 effector molecules known to either activate or inhibit transcription factors (extracted from (Delépine et al., 2016)). SEMPs were generated by enumerating pathways linking source to sink in a single iteration by firing rules computed from MetaNetX (provided in supplementary materials).

Among the 421 biomarkers, we found 27 biomarkers directly detectable by transcription factors, and 415 pathways enabling the transformations of 164 different biomarkers into 76 different effectors. Some of these results are presented in Table 2.

**Table 2.**
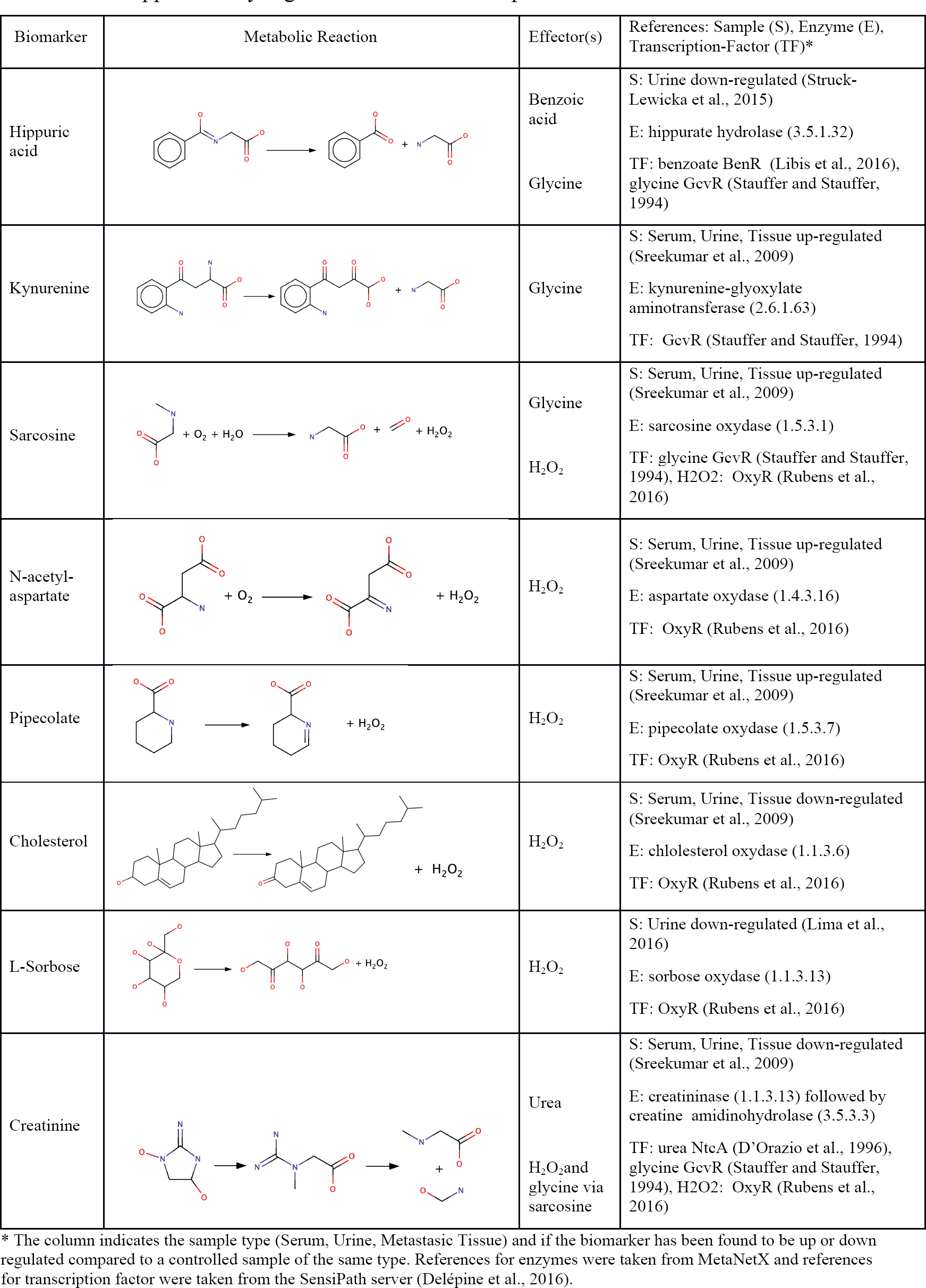
Examples of metabolic pathways enabling the detection of prostate cancer biomarkers. Supplementary Figure 2 illustrates the scope of sarcosine.

| Biomarker | Metabolic Reaction | Effector(s) | References: Sample (S), Enzyme (E), Transcription-Factor (TF)* |
| --- | --- | --- | --- |
| Hippuric acid |  | Benzoic acid<br>Glycine | S: Urine down-regulated (Struck-Lewicka et al., 2015)<br>E: hippurate hydrolase (3.5.1.32)<br>TF: benzoate BenR (Libis et al., 2016), glycine GcvR (Stauffer and Stauffer, 1994) |
| Kynurenine |  | Glycine | S: Serum, Urine, Tissue up-regulated (Sreekumar et al., 2009)<br>E: kynurenine-glyoxylate aminotransferase (2.6.1.63)<br>TF: GcvR (Stauffer and Stauffer, 1994) |
| Sarcosine |  | Glycine<br>H<sub>2</sub>O<sub>2</sub> | S: Serum, Urine, Tissue up-regulated (Sreekumar et al., 2009)<br>E: sarcosine oxydase (1.5.3.1)<br>TF: glycine GcvR (Stauffer and Stauffer, 1994), H2O2: OxyR (Rubens et al., 2016) |
| N-acetyl-aspartate |  | H<sub>2</sub>O<sub>2</sub> | S: Serum, Urine, Tissue up-regulated (Sreekumar et al., 2009)<br>E: aspartate oxydase (1.4.3.16)<br>TF: OxyR (Rubens et al., 2016) |
| Pipecolate |  | H<sub>2</sub>O<sub>2</sub> | S: Serum, Urine, Tissue up-regulated (Sreekumar et al., 2009)<br>E: pipecolate oxydase (1.5.3.7)<br>TF: OxyR (Rubens et al., 2016) |
| Cholesterol |  | H<sub>2</sub>O<sub>2</sub> | S: Serum, Urine, Tissue down-regulated (Sreekumar et al., 2009)<br>E: cholesterol oxydase (1.1.3.6)<br>TF: OxyR (Rubens et al., 2016) |
| L-Sorbose |  | H<sub>2</sub>O<sub>2</sub> | S: Urine down-regulated (Lima et al., 2016)<br>E: sorbose oxydase (1.1.3.13)<br>TF: OxyR (Rubens et al., 2016) |
| Creatinine |  | Urea<br>H<sub>2</sub>O<sub>2</sub> and glycine via sarcosine | S: Serum, Urine, Tissue down-regulated (Sreekumar et al., 2009)<br>E: creatininase (1.1.3.13) followed by creatine amidinohydrolase (3.5.3.3)<br>TF: urea NtcA (D'Orazio et al., 1996), glycine GcvR (Stauffer and Stauffer, 1994), H2O2: OxyR (Rubens et al., 2016) |

\* The column indicates the sample type (Serum, Urine, Metastatic Tissue) and if the biomarker has been found to be up or down regulated compared to a controlled sample of the same type. References for enzymes were taken from MetaNetX and references for transcription factor were taken from the SensiPath server (Delépine et al., 2016).

Notable amongst the biosensor listed in Table 2 are H_2_O_2_ and glycine that are detectable by the native *E. coli* transcription factors OxyR and GcvR, respectively (Tartaglia et al., 1989; Stauffer and Stauffer, 1994), and benzoate for which biosensors have already been built in *E. coli* (Libis et al., 2016) (detailed results are provided in the supplementary materials). Interestingly several biomarkers could be transformed into the same effector, thus enabling the integration of multiple biomarker signals into a unique detectable biosensor.

Those results highlight the versatile use that a generic retrosynthesis and reaction network prediction algorithm such as RetroPath2.0 can have.

## 5. Discussion

The RetroPath2.0 workflow is a versatile reaction network tool, built to be modular enough to answer most metabolic engineering needs. RetroPath2.0 takes as input a first set of compounds (the source), a second set of compounds (the sink) and a set of reaction rules (see Figure 3). The workflow produces a network linking the source set to the sink set, where each link in the network correspond to a reaction rule. The RetroPath2.0 workflow runs under the KNIME analytics platform and it is available in Supplementary Material at https://www.myexperiment.org/workflows/4987.html.

The choice of source, sink and rule sets depends on the application. For instance, if one wishes to find all possible synthesis routes that can be engineered for a target compound, the source set is the target, the sink is the set of metabolite of the chassis strain, and the rules are the reversed form of all known metabolic reactions (cf. 4.2.1). If one is interested in finding pathways to be engineered to degrade a given xenobiotic, the source set is the xenobiotic, the sink set can be composed of those metabolites in the central metabolism of a chassis strain and the rule set could comprise all known catabolic reactions. In the same vein, one can find sensing-enabling pathways with the set of known detectable compounds as sink, the set of target compounds to detect as source, and using forward rules (cf. 4.2.2 for the detection of biomarkers). Finally if one wishes to know all possible compounds that can be produced with a chassis strain when adding heterologous enzymes, the source set is composed of the metabolites of the chassis strain, the sink set can be either empty or a set of compounds in a vendor catalogue, and the rule set should cover all reactions that could occur in the chassis strain, including heterologous enzymatic ones. Moreover, any other applications where the problem can be framed into source, sink and rule sets can be processed by the workflow including problems where compounds are not metabolites and reactions are not metabolic reactions.

The most critical feature of a reaction network prediction system is certainly how the reactions are encoded and from where this knowledge was extracted. In our case, we choose to adopt a reaction encoding based on SMARTS, a widely accepted compound query language (Daylight, 2017) that was already used successfully in such context (Hadadi and Hatzimanikatis, 2015). Unlike most rule-based reaction prediction systems, RetroPath2.0 rules are not built around the Enzyme Commission nomenclature, but rather from an automatic translation of enzymatic reactions extracted from databases, which we believe offers a refined view of enzyme’s capabilities.

We showed that our rules were able to classify reactions and that our set of rules extracted from MetaNetX had a good coverage over the known reactome. A good part of the reactions that were not covered were actually reactions involving compound classes (e.g. “an alcohol”), which were removed during the rule generation steps. This type of generalized reactions were, in turn, represented in our set through our unique way of encoding reactions as generalized rules. One substantial improvement could probably be met by constraining the atom-atom mapping and reaction centre identification steps based on the exploitation of additional knowledge on the reaction and the associated enzyme. For instance by using the known alternative substrates associated to a single enzyme sequence, or the EC number assignation.

Evaluating the coverage of a reaction database is interesting in order to assert the coverage of the known reactome by a given set of rules, but it cannot be used to assert the efficiency of a retrosynthesis tool. Indeed, the coverage of a reaction database depends mainly on the database from which the rules were inferred and how exhaustive the cross-links are between those two. Ideally, we would want a set of rules to be able to recover all known biochemical reactions. It means that anything less than 100% coverage is an admission that the set of rules is incomplete and that more data could have been aggregated. Note that in this work we focus our efforts on MetaNetX for the sake of simplicity but it is clear that more data can be imported from other databases such as BRENDA (Chang et al., 2015).

To the author's opinion, a better indicator of retrosynthesis tools efficiency should be found in the coverage of known pathways realized in a metabolic engineering context. This is precisely what we did using the LASER database as a reference for examples of successfully engineered metabolic pathways. In that way we provided a comprehensive overview of the capabilities of our tool in order to identify metabolic engineering solutions to bioproduction for well-studied cases. The main source of misprediction that we observed in our analysis came from cases in which additional compounds absent from *E. coli* metabolism were needed to perform the synthesis. Indeed, we performed all our computations within five iterations from *E. coli,* with target compounds that were not necessarily produced in this chassis, nor at five enzymatic steps; moreover, some substrates could be supplemented in the media of the chassis organism. For instance, the synthesis of morphine is described for *Saccharomyces cerevisiae* in (Thodey et al., 2014) by two pathways at three and four steps from thebaine. Thebaine is not naturally present in *E. coli* metabolism thus absent from the sink we used. Consequently, this example has no scope at five steps and was counted as mispredicted. Once thebaine is supplemented in the sink, RetroPath2.0 can generate a scope with both pathways. Note that thebaine was already predicted before being added to the sink, and that doing so only allowed RetroPath2.0 to use this compound as a valid starting point for synthesis instead of continuing further the retrosynthesis.

Importantly, not all predicted pathways can readily be implemented in *E. coli.* Indeed, translation of *in silico* models into *in vivo* experiments require much more constraints to be satisfied, some of those being hardly predictable. To name but a few, enzyme sequence availability, chassis ability to fold the enzymes, kinetics, intermediate compounds toxicity, and overall pathway induced stress on the cell should all be checked before going any further. In this context, RetroPath2.0 can be seen as a base on which everyone is invited to build new features in order to further improve its metabolic space exploration abilities.

Exploiting chemical diversity in order to gain access to the large catalogue of natural and non-natural chemical resources is arguably one of the most important goals of biotechnology applications. By extending metabolic capabilities of enzymes, applications in metabolic engineering, biosensors and synthetic circuits can be greatly enlarged and diversified. To that end, RetroPath2.0 brings to the community a flexible and scalable open source platform with unique metabolic design capabilities. For the first time, we allow the systematic application of a full set of validated and standardized reaction rules that can be expressed with a selectable level of specificity. Such representation, which parallels the versatility of enzyme promiscuity, allows an in-depth exploration of latent abilities of natural enzymes.

The excellent coverage of the workflow along with its proved ability for recovering both known pathways and putative alternative candidate pathways show its power as an engineering tool. For that reason, we have no doubt that the tool will be received as a valuable addition to the toolbox for engineering biology. Moreover, community contributions to the workflow will likely expand further the features of the tool, even beyond metabolic design. In summary, we believe that the ability of RetroPath2.0 to rationalize and standardize design steps of biological engineering that have been traditionally performed manually by trial and error, constitutes a major contribution towards the development of automated workflows across the whole design, build, test and learn cycle.

